# From Spectra to Digital Phenotypes: Wearable Multispectral Sensing for Precision Light and Green Space Exposure

**DOI:** 10.64898/2026.05.14.724799

**Authors:** Rong Liu, Yuchong Han, Huayi Lu, Yifeng Zhou, Tian Xue

**Affiliations:** Department of Ophthalmology, The First Affiliated Hospital of USTC, State Key Laboratory of Eye Health, New Cornerstone Science Laboratory, Biomedical Sciences and Health Laboratory of Anhui Province, School of Life Sciences, Division of Life Sciences and Medicine, University of Science and Technology of China, Hefei 230026, China

## Abstract

Light is a modifiable determinant of health, yet real-world exposure assessment is often reduced to illuminance alone, lacks environmental context, or relies on privacy-sensitive sensing. We present SpectraVita, a low-cost, compact multispectral wearable that continuously samples 11 ultraviolet-to-near-infrared bands and, through a privacy-preserving pipeline without cameras or location tracking, produces interpretable digital phenotypes of lighting environment (natural vs. artificial and source type) and vegetation context alongside standard visual and non-visual light metrics. In extensive in-the-wild recordings spanning diverse scenes, times of day, weather conditions, and light sources, we observe distinctive spectral signatures that enable supervised models to achieve a macro-averaged F1 score of 0.988±0.004 for light-source classification and green-space detection in boundary-free environments. A sensor-derived normalized difference vegetation index (NDVI) emerges as an explainable, physically grounded marker linking natural light exposure and greenness. Robustness is supported by scenario-shift testing, image-segmentation validation, and mixed-environment experiments that demonstrate sensitivity to partial and transient exposures, as well as by longitudinal stationary monitoring and deployment in a cohort of thousands of participants capturing seasonal and behavioral variability. SpectraVita enables individualized, privacy-preserving, longitudinal monitoring of light and greenness exposure at scale, addressing a key measurement gap for precision and population health studies of daily photic environments.

## Main Text

Light exposure is a key modifiable factor influencing sleep, circadian timing, mood, ocular health, cardiometabolic function, and cognition across the lifespan(see reviews^1–9^). Emergent evidence indicates that appropriate natural light exposure supports mental and physical well-being^10–13^, whereas artificial light, particularly at night, can disrupt circadian regulation and physiological homeostasis^14–18^. However, the health effects of light are not homogeneous: they depend critically on spectral composition, timing, and environmental context. Among natural light environments, exposure near a window, outdoors in open sky, or under a vegetated canopy can yield meaningfully different spectra and intensities, with potentially different implications for vitamin D synthesis, eyesight, and visual comfort^13, 19^. In particular, green space exposure—often operationalized via satellite-based vegetation indices—has been linked to better mental health, and lower all-cause mortality^20–25^. Among artificial sources, for example, fluorescent lamps has been associated with eye strain and headaches^26^. Translating these nuanced insights into actionable digital health monitoring and personalized interventions requires continuous, individual-level characterization of the lived light environment, including both source type (e.g., LED, fluorescent, incandescent, daylight) and context (indoor, outdoor open, green space), a capability that remains limited in current precision medicine.

Existing measurement approaches remain fundamentally limited for this purpose^27, 28^. Conventional wearable illuminance meters capture overall intensity but ignore spectral composition and the contextual structure of the light field. Devices that reconstruct spectra^29, 30^ or report α –opic equivalent daylight illuminance^31–33^ improve physiological relevance, but typically still do not identify the dominant source type or the environmental setting in which exposure occurs. Satellite-based measures provide coarse, location-based proxies of outdoor lighting and green space^34, 35^ but cannot capture personal exposure trajectories or the spectrum reaching the eye. Wearable cameras can supply human-centric context and have been used to estimate green space exposure^36^, but raise substantial privacy concerns and impose weight, computational, and cost burdens that limit acceptability, especially at population scale.

Here we introduce SpectraVita, a compact wearable multispectral detector that continuously samples ultraviolet, visible, and near-infrared light across 11 channels. Coupled with a privacy-preserving pipeline that avoids imaging and location tracking, SpectraVita computes standard visual and non-visual metrics (e.g., illuminance, correlated color temperature, α-opic equivalent daylight illuminance) and generates interpretable digital phenotypes of lighting source and environmental context. We further establish that a vegetation index derived from multispectral channels (NDVI) serves as a sensitive, explainable marker for both natural light and green space exposure. Large-scale real-world deployments spanning thousands of participants demonstrate robustness across seasons and behavior, supporting scalable, context-aware inference of individual light exposure with minimal privacy risk. SpectraVita therefore enables practical digital health applications—from circadian-informed monitoring and personalized light-based interventions to precision medicine studies requiring continuous measurement of both light and green space exposure.

## Results

### SpectraVita workflow

Figure 1 summarizes the SpectraVita workflow, designed to translate raw wearable multispectral measurements into interpretable, digital phenotypes of real-world lighting context for digital health monitoring. The pipeline is organized into three modules: (1) on-body acquisition of multi-channel ultraviolet/visible/near-infrared signals; (2) data preprocessing; (3) feature generation through a dual-path strategy that yields both conventional light metrics and innovative lighting environment and vegetation context.

**Fig. 1.**
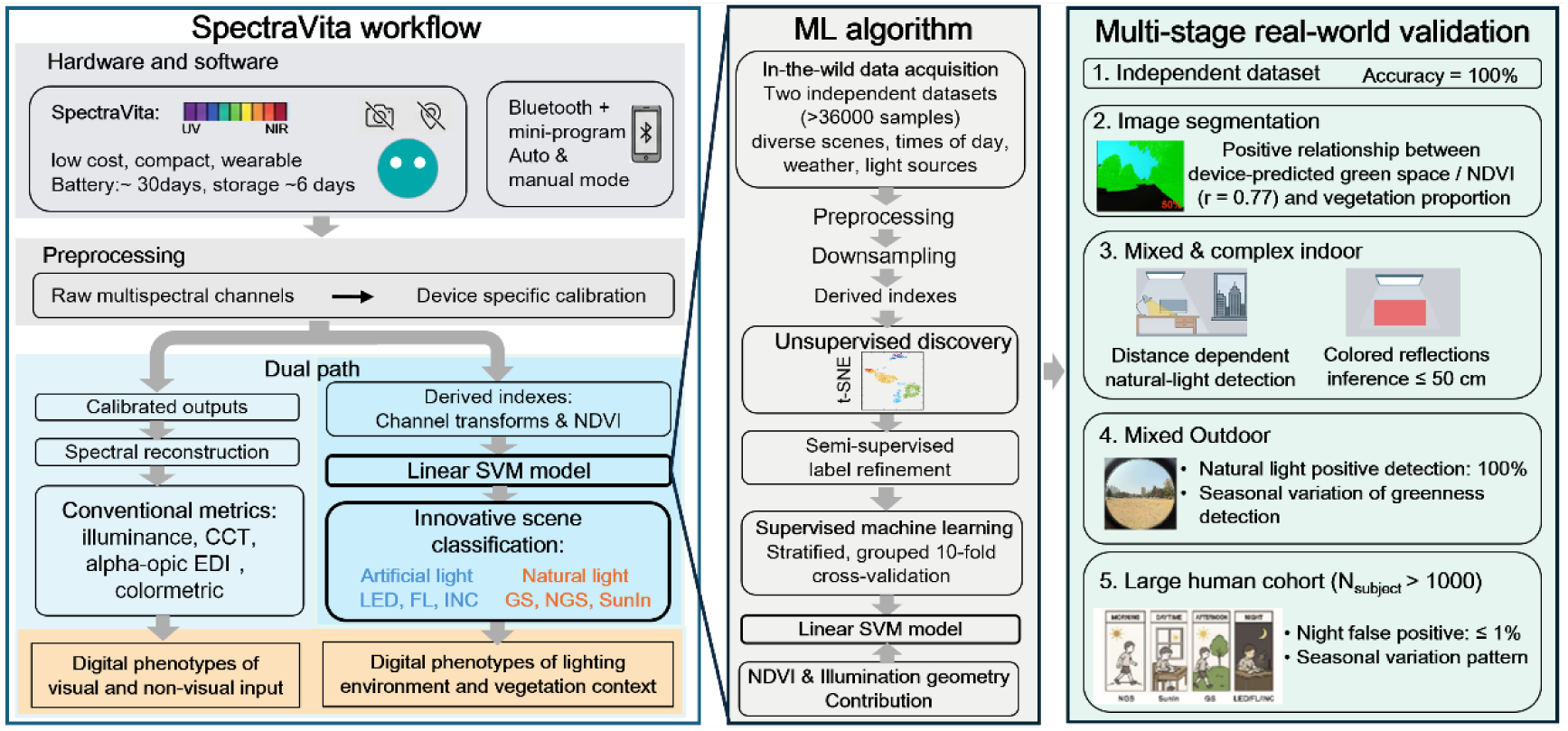
Schematic illustration of the SpectraVita workflow, including the machine learning framework (middle panel) and multi-state real-world validation (right panel). The workflow comprises three modules: (1) on-body acquisition of multi-channel ultraviolet, visible, and near-infrared signals; (2) data preprocessing; and (3) feature generation via a dual-path strategy that yields both conventional light metrics and novel descriptors of lighting environment and vegetation context. A linear SVM model for classifying lighting environment and vegetation context was trained and validated on in-the-wild recordings spanning diverse scenes, and further evaluated in a large human cohort.

SpectraVita—a compact wearable multispectral detector captures full-spectrum light across 11 channels spanning UV, visible, and NIR bands. Importantly, the system is intentionally camera-free and does not require GPS, enabling context inference with substantially reduced privacy risk compared with vision– or location-based approaches. In routine operation, the device records continuously at fixed intervals (5 min in the default “auto” mode), a configuration selected to balance temporal resolution, battery life, and storage requirements; users upload data via Bluetooth using a companion mobile mini-program that also provides real-time visualization. A “manual” mode supports targeted testing under specific environments by allowing dynamic gain adjustment to extend the measurable intensity range and by enabling an additional flicker-detection channel (see Methods and Supplementary Materials).

Following acquisition, raw channel outputs are processed with device-level calibration coefficients to ensure cross-device comparability and to support reconstruction of spectral quantities. The pipeline then follows a dual-path feature strategy. In the conventional path, reconstructed spectra are used to compute standard photometric and circadian-relevant metrics (e.g., illuminance, correlated color temperature, and CIE S026 alpha-opic equivalent daylight illuminance), which facilitates comparability with existing spectrometer. These metrics are validated against a reference spectrometer, achieving high linearity (R² > 0.99) and accuracy (see Methods and Supplementary Materials). In parallel, Innovative machine learning path derives spectrum-centric features, including channel transformations and the normalized difference vegetation index (NDVI). Unsupervised discovery reveals intrinsic spectral structure, guiding semi-supervised label refinement. A linear support vector machine (SVM) classifier, trained with 10-fold cross-validation and SMOTE (the Synthetic Minority Over-sampling Technique) balancing, then predict six environmental categories: green space (GS), non-green space (NGS), and sunlit indoor (SunIn), LED, fluorescent (FL), incandescent (INC), These outputs form digital phenotypes of lighting environment and vegetation context. Together, this integrated framework establishes SpectraVita as a scalable, privacy-compliant tool for generating clinically actionable digital phenotypes of visual and non-visual input, and individual light and green space exposure.

### Unsupervised discovery reveals intrinsic spectral structure

To evaluate whether wearable multispectral measurements capture meaningful environmental variation in real-world settings, we collected two independent datasets using SpectraVita: a primary dataset model development (training and cross-validation), and a separate validation dataset acquired under completely different scenarios to assess model generalizability. Data encompassed six representative real-world lighting environments: including LED, fluorescent light (abbreviated as FL), incandescent light (INC), sunlit indoor (SunIn), green space (GS), and non-green space (NGS) conditions)—covering both artificial and natural contexts. Measurements were collected across heterogeneous indoor and outdoor settings (e.g., streets, schools, residential communities, parks, playgrounds, laboratories, hospitals, kindergartens, hotels, and homes), over the full daytime cycle (sunrise to sunset), under varied weather (sunny, cloudy, rainy), and across seasons (winter to summer). Representative examples are shown in Fig. 2a.

**Fig. 2.**
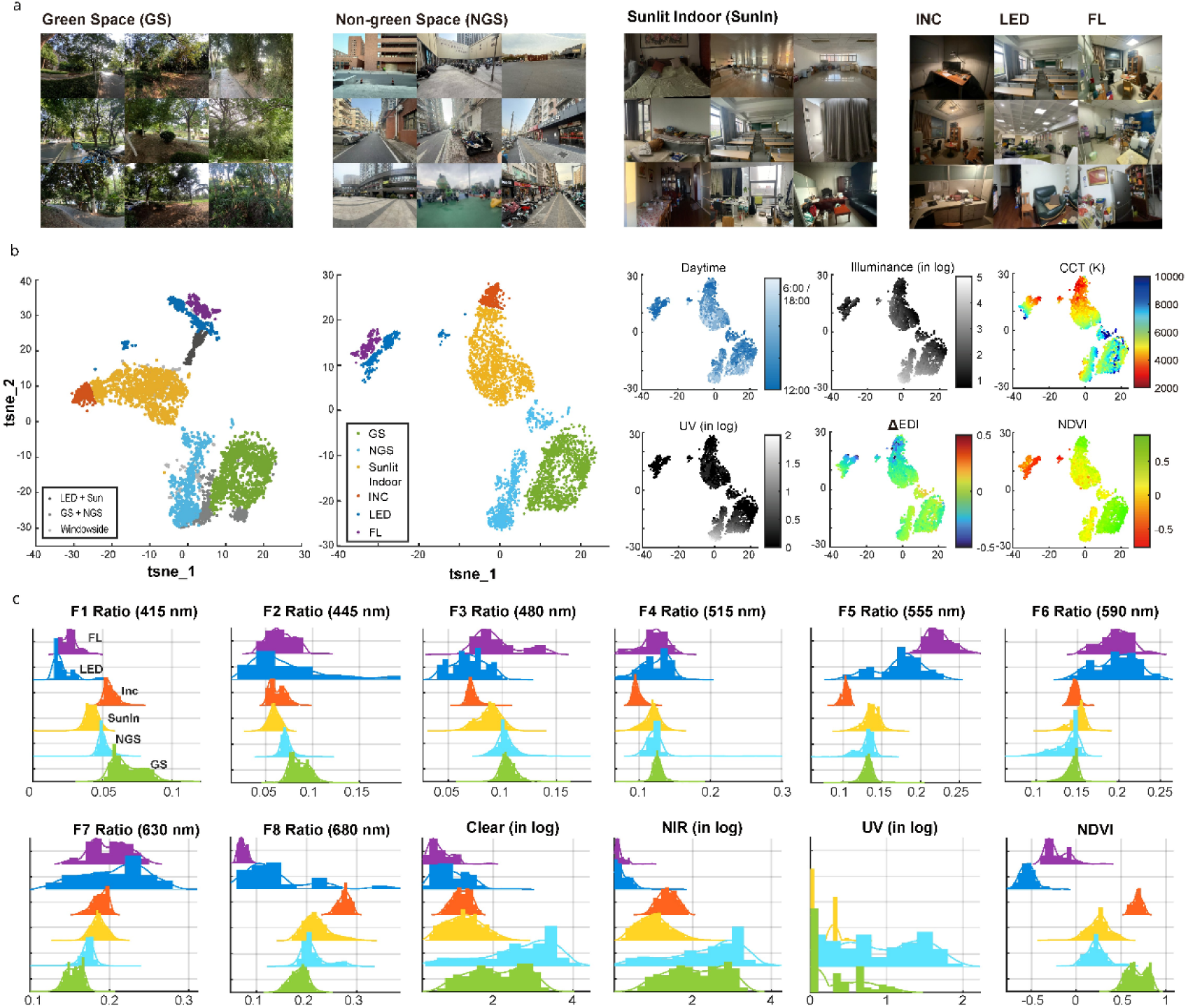
a) Example photographs of six representative real-world lighting environments from which data were collected. b) t-SNE analysis. The left panel shows clear segregation of the six major lighting environments categories, with intermediate mixed-lighting conditions located between clusters. The middle panel shows the final dataset after removal of mixed-lighting records. The smaller panels on the right illustrate representative feature distributions, with individual samples color-coded and annotated by collection time or light metrics. c) The distribution of the outputs of each channel and the derived NDVI values across different light sources. The solid line represents the normalized probability density function. The peak wavelength of each visible channel was shown in parentheses. F1-F8 channels were scaled to the sum of their total outputs. Clear, NIR and UV channels were original outputs in logarithm. NDVI was computed as (Nir-F8)/ (Nir+F8), using the raw NIR and F8 signals.

Raw sensor signals underwent systematic preprocessing. Visible channels (F1–F8) were normalized by their total output to enable channel-independent comparisons and reduce inter-device variability, achieving a median coefficient of variation of 1% ± 1% across visible channels and illuminance levels. Clear, NIR (near Infrared), and UV (ultraviolet) channels were log-transformed. NDVI, a standard indicator of greenness, was calculated as (NIR − F8)/(NIR + F8) using original sensor outputs. Samples with derived illuminance ≤ 5 lux were excluded from analysis.

We first applied t-distributed Stochastic Neighbor Embedding (t-SNE) to the combined datasets to visualize intrinsic spectral relationships without imposing labeled boundaries. The resulting embedding revealed well-separated clusters aligned with major lighting contexts, indicating that the multispectral signatures contain strong intrinsic structure (Fig. 2b, first panel). Importantly, mixed lighting scenarios (e.g., meadow edges) formed intermediate manifolds between cluster centers rather than collapsing into a single category, consistent with continuous mixing of spectral components. Similarly, SunIn measurements near windows appeared across both SunIn and NGS regions, reflecting hybrid illumination with contributions from both indoor and outdoor sources. Because some labeled categories (notably GS/NGS and SunIn/NGS) are intrinsically ambiguous in real-world conditions, we used the t-SNE structure as a data-driven reference to refine labels for subsequent supervised analyses. Specifically, we semi-supervisedly curated GS/NGS and SunIn/NGS by excluding clear outliers falling outside their corresponding cluster regions. After removing strongly mixed-light records, we obtained a final dataset comprising six well-defined categories (Fig. 2b, second panel), providing a cleaner basis for classification and for later analyses of boundary cases.

Feature exploration highlighted NDVI as a sensitive and physically interpretable index. NDVI effectively discriminating both GS vs. NGS conditions, and cool vs. warm light sources—surpassing conventional metrics like intensity and correlated color temperature (CCT) (Fig. 2b, right panels). Histogram analyses of sensor channel responses and derived lighting indices (Fig. 2c) further illustrated the pronounced variation in NDVI across lighting conditions, providing additional discriminative power for natural vs. artificial illumination and GS vs. NGS. These exploratory findings motivated inclusion of NDVI alongside raw channels in the supervised feature set.

### Supervised light-context classification with independent validation

Following quality control, the datasets were separated based on their original collection sources into a primary dataset (4,309 samples) and an independent validation dataset (293 samples, see details in supplementary table 1-2). Multiple supervised machine-learning algorithms were evaluated on the primary dataset using stratified, grouped 10-fold cross-validation to classify lighting environments. To address class imbalance among sub-categories, the Synthetic Minority Over-sampling Technique (SMOTE) was applied until all sub-categories contained comparable numbers of observations.

A linear Support Vector Machine (SVM) achieved the best performance, with a macro-averaged F1 score of 0.988±0.004 and near-perfect accuracy (>99.9%) for coarse classification tasks such as natural vs. artificial lighting and cool vs. warm sources (natural lighting and incandescent light) (Fig. 3a). When evaluated on the independent scenario-shift validation dataset collected under distinct scenarios, the trained model maintained 100% accuracy, supporting robust generalization across heterogeneous real-world scenes.

**Fig. 3.**
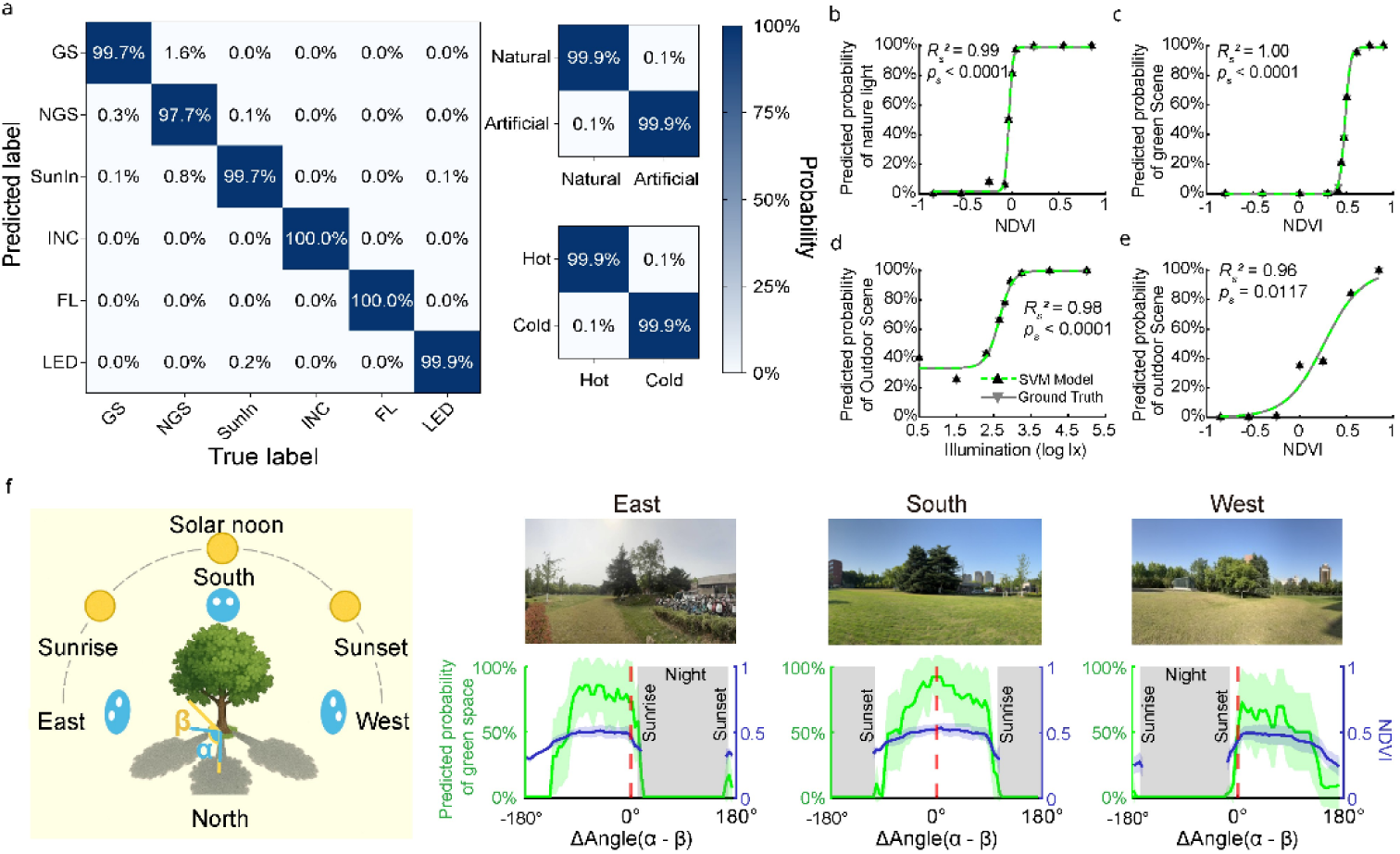
a) Prediction performance of the SVM model on the primary dataset. b) Relationship between NDVI and predicted/true proportion of natural light. c) Relationship between NDVI and predicted/true probability of green space (GS). d) Relationship between illumination and predicted/true probability of outdoor scenes. e) Relationship between NDVI and predicted/true proportion of outdoor scenes. f) Left: Schematic of illumination-geometry modulation. Green-space predictions and NDVI are highest when the device azimuth (α) and solar azimuth (β) are aligned with the target vegetation (i.e., α − β = 0; device viewing direction and solar illumination direction both point toward the same vegetated target). Right: Relationship between green-space predictions/NDVI and the angular difference α and β during real-world long-term monitoring. Solid green lines in the figures indicate the across-day mean, and shaded areas denote 95% confidence intervals. Red dashed lines indicates α – β = 0. The photograph above each curve illustrates the corresponding sensor view.

Analysis of linear SVM coefficients across cross-validation folds revealed NDVI as the dominant predictor, with mean absolute β-weight approximately 1.7-fold higher than the second most influential variable, ranking first in every fold. We then related NDVI directly to model outputs for two key decisions: natural versus artificial light and GS versus NGS. Across the two combined datasets (excluding incandescent light, which is artificial but exhibits high NDVI), NDVI showed a clear sigmoidal relationship with the model’s predicted probability of natural light (Fig. 3b): samples with NDVI > 0 were consistently classified as natural light, with a half-saturation point at −0.04, indicating that NDVI captures spectral composition in a way that is robust to variation in absolute brightness. Similarly, green space classification probability increased sigmoidally with NDVI (half-saturation: 0.49, Fig. 3c).

Previous studies often used an illuminance cutoff of 1000 lux^37, 38^ as a threshold to distinguish outdoor from indoor conditions. In our dataset, when samples were binned by log illuminance, the empirical fraction of outdoor labels (GS+NGS) increased with illuminance, with a 50% predicted probability at 285 lux; at 1,000 lux, the outdoor probability reached 92% (Fig. 3d). However, a substantial fraction of outdoor exposures—particularly GS—occurred below 1,000 lux in our data. Therefore, an illuminance-only cutoff would systematically miss low-illuminance outdoor exposures (false negatives). In contrast, incorporating spectral phenotypes such as NDVI (Fig. 3e) enabled accurate classification across a wide illuminance range and in spectrally complex environments.

### Illumination geometry modulates NDVI and detection sensitivity

Further analysis showed that both NDVI and model’s green-space probability varied systematically with the angular difference between the device azimuth and the solar azimuth relative to the target vegetation (Fig. 3f). Green-space probabilities were highest when the two azimuths were closely aligned, indicating that directional illumination geometry modulates vegetation reflectance and, in turn, influences the classifier output. This pattern suggests that our detection is sensitive to illumination-dependent spectral cues rather than relying solely on image-based “greenness”.

### Image-segmentation ground-truth validation of green-space detection

As an independent verification vegetation detection, the multispectral sensors were paired with a wide-angle action camera (GoPro HERO13, GoPro, Inc., US) to synchronously capture spectral indices and corresponding imagery. Semantic segmentation of the images provided ground-truth vegetation proportions (Fig. 4a), which correlated linearly with NDVI (Fig. 4b, r = 0.77 *p* < 0.0001, Pearson correlation, N = 320) and exhibited sigmoidal relationship with green-space prediction probability (Fig. 4c, grey line). When vegetation coverage exceeded 53%, green-space probability rose above 50% and plateauing near 88%. Notably, probabilities approached 100% only under tree canopy (Fig. 4c, green line), reflecting NDVI’s dependence on vegetation type and canopy structure^39^—a nuance that enhances rather than compromises the biomarker’s interpretability.

**Fig. 4.**
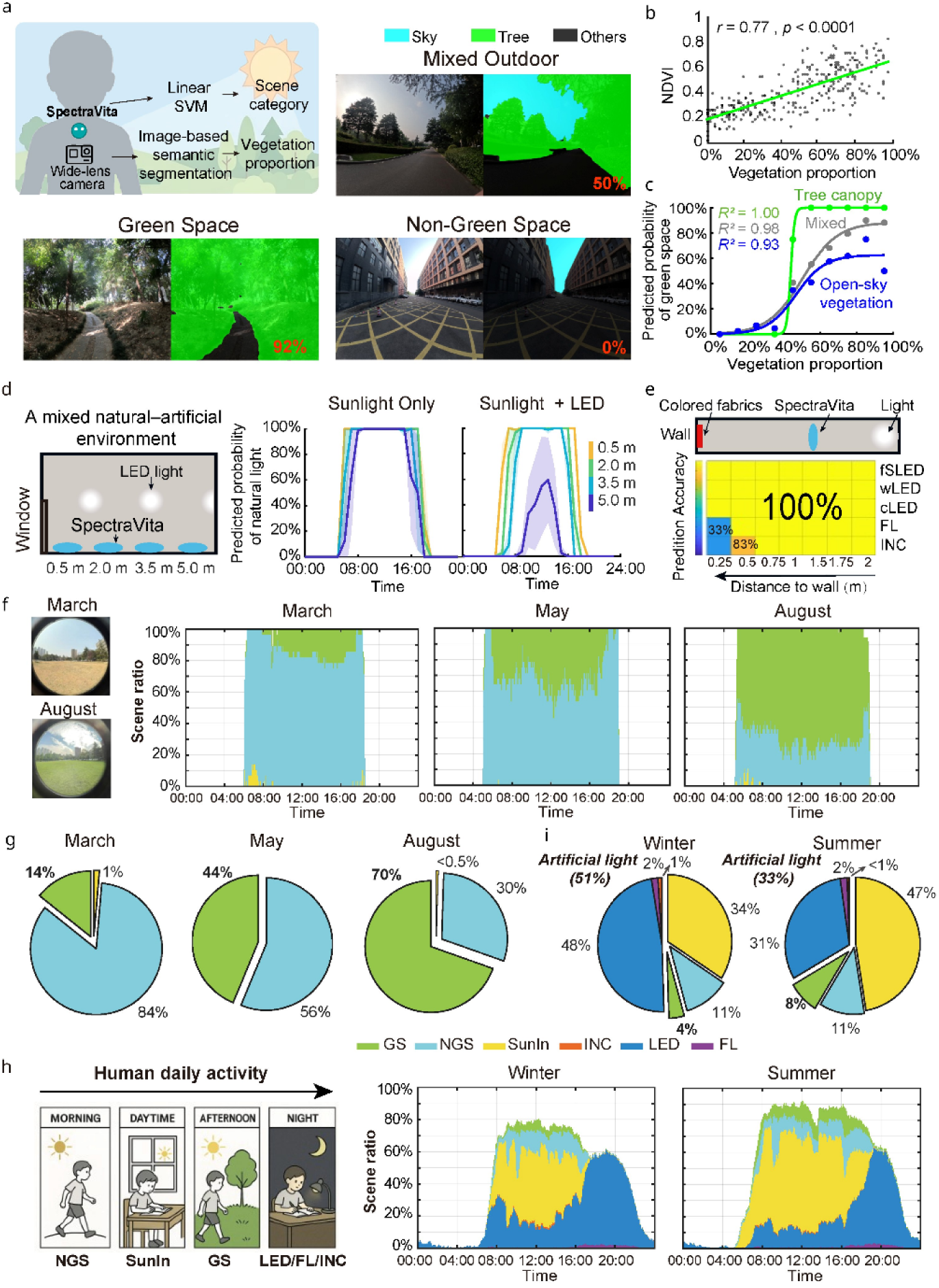
a) Top left: schematic of the image-segmentation validation. Remaining panels: example paired images showing semantic segmentation applied to representative outdoor scenes; pixels labeled “sky” and “tree” are colored blue and green, respectively. The vegetation proportion (fraction of pixels labeled “tree”) is indicated in the lower-right of each image. b) Correlation between vegetation proportion (semantic segmentation) and NDVI measured by SpectraVita. c) Relationship between vegetation proportion and the predicted probability of green space from SpectraVita. Curves are shown separately for observations collected under tree canopy (green), open-sky vegetation (blue), and mixed conditions (grey). d) Left: top-view schematic of an indoor space jointly lit by daylight and LED. Right, model predictions of natural-light probability over time in mixed indoor environments. Solid lines show across-day means, and shaded areas denote 95% confidence intervals. e) Heat map showing that prediction accuracy increases with greater distance between reflective colored fabric and the sensor. Top: top-view schematic of the indoor test with brightly colored fabrics placed on a wall. Abbreviations: fsLED = full-spectrum LED; wLED = warm LED; cLED = cold LED; FL = fluorescent; INC = incandescent. (f) Left: fish-eye view from one of four sensors, shown across seasons. Right, stacked scene categories across an average 24 h day for open meadow measurements in different seasons. (f) Pie charts of the grand average daily proportion of light-exposure categories for open meadow in different seasons. h) Left, schematic illustration of a typical day for participating children (partly AI-generated). Right, stacked scene categories across an average 24 h day, shown separately for each season. Colors denote lighting scenes; blank segments indicate time points excluded due to non-wear or sensor occlusion. On average, ≈15% of daytime data were excluded, with a higher proportion at night during sleep. The midday dip reflects common nap habits among Chinese schoolchildren. i) Pie charts showing grand average daily proportion of light-exposure categories for human participants, separately for each season.

### Operational robustness under mixed lighting and reflective interference indoor environments

We evaluated robustness to mixed illumination and proximal colored reflections. In indoor space jointly lit by daylight and LED, the predicted probability of “sunlight” decreased systematically with increasing distance from the window—consistent with the spatial gradient of natural-light contribution. Conversely, when artificial lighting was switched off during daytime, the model accurately identified sunlight (Fig. 4d). Despite the high accuracy of natural-light detection, measurements taken near windows were frequently labeled as “NGS” (24.2% on average at 0.5 m from the window, decreasing to 0% at larger distances) rather than “SunIn”, reflecting the fact that the wearable captures the spectral composition of the light field rather than the physical “indoor/outdoor” location. This behavior is consistent with the mixed manifolds observed in the t-SNE embedding and highlights boundary conditions where hybrid illumination is expected, and is consistent with previous work using light-based metrics to classify indoor versus outdoor environments^40^.

Potential interference from colored reflections was tested using brightly colored fabrics placed on walls under varied lighting conditions (three LED lights with different CCTs, an incandescent light, and a fluorescent light). Measurements at different distances between sensor and surface showed potential interference from colored reflections occurred only at distances ≤50 cm (at which point colored surfaces dominated the sensor’s 66 × 84° field of view, Fig. 4e), substantially closer than typical wearable use. This suggests that, in normal operation, the device predominantly captures ambient illumination rather than proximal reflected color spectra.

### Validation in real-world long-term stationary outdoor monitoring

We validated the SVM-based lighting classifier in real-world long term stationary monitoring. In untrained mixed outdoor settings (open meadow in USTC campus, Hefei, China) where illumination should be purely natural, the model predicted natural light with 100% accuracy across three seasonal sessions.

The model’s green-space predictions tracked seasonal vegetation changes, increasing from 14% GS / 84% NGS / 1% SunIn in March (total of 9 days monitoring from 4 devices each) to 44% GS / 56% NGS in May (7 days) and 70% GS / 30% NGS in August (7 days), demonstrating sensitivity to greenness changes over time (Fig. 4f-g).

### Validation in real-world large cohort study

We further validated the SVM-based lighting classifier in real-world deployments to assess performance under true wear conditions. We examined night-time data (20:00–04:00, illuminance >10 lux), when illumination is expected to be artificial, across a large cohort of school-aged children. Among 218,158 samples from 738 participants collected in Oct–Dec 2023, the model incorrectly labeled only 0.7% of samples as natural light. Similar rates were observed in May–June 2024 (0.6% of 147,260 samples from 570 participants) and Dec 2024–Jan 2025 (1.0% of 199,502 samples from 802 participants), indicating a nighttime false natural-light detection rate of ≤1% across seasons.

Longitudinal summaries further revealed clear seasonal structure in individual exposure patterns, (including 193 children who participated in both May–June 2024 and December 2024–January 2025). In early summer participants experienced 67 ± 9% of eligible time under natural light and 33 ± 9% under artificial light, with LED sources comprising 31 ± 10% of total time (≈ 3.4 h/day). Non-green-space (NGS) exposure averaged 11 ± 4% (≈ 1.4 h/day) and green-space (GS) exposure 8 ± 10% (≈ 1.0 h/day). In winter, natural light fell to 49 ± 10% while artificial light rose to 51 ± 10%; LED contribution rose to 48 ± 11%, NGS remained stable at 11 ± 3% (≈ 1.2 h/day), whereas GS exposure declined to 4 ± 3% (≈ 0.4 h/day) (Fig. 4h-i). Collectively, these large-scale real-world results support the classifier’s in situ accuracy and demonstrate sensitivity to seasonal and behavioral variation in both lighting and greenness exposure.

## Discussion

This study demonstrates that wearable multispectral sensing combined with a privacy-preserving computational pipeline enables accurate, scalable, and interpretable characterization of individual light exposure (natural vs. artificial light, green space vs. non-green space.) in real-world settings—addressing a key need in precision environmental health. SpectraVita generates conventional visual and non-visual metrics as well as contextual digital phenotypes, including light-source identification and green-space detection, without cameras or GPS. This spectral–context fusion extends recent spectrally aware loggers and improves on conventional broadband and single-band devices (See reviews^41, 42^) which largely neglect ultraviolet or near-infrared domains. By achieving robust phenotyping with substantially reduced privacy risk, the system overcomes a critical barrier to widespread adoption in population-scale studies.

Accurate discrimination between natural and artificial light sources is essential for understanding how built environments reshape human photic input. Early studies typically relied on survey of indoor and outdoor activities^43, 44^, which are not objective and lack time resolution. More recent approaches using illuminance thresholds^38^ or UV intensity^45^ struggles in modern societies where individuals spend most time indoors under mixed lighting. Our multispectral system achieves near-perfect separation (>99% accuracy) in clear-boundary environments while remaining sensitive to spectral gradients in mixed settings, enabling objective light-source annotation without reliance on time or location heuristics.

Green space exposure is widely recognized as a determinant of health^20–24^, with benefits spanning mental well-being, cognitive function, and mortality. Traditional quantification typically links residential addres^34, 46^ and Global Positioning Systems (GPS)^47^ or Location-Based Services (LBS)^48^ trajectory to satellite-derived greenness indices such as NDVI^49–51^. Streetscape greenery has also been assessed using street-view data^35, 52^ or wearable/smartphone cameras^36, 53^ with image segmentation. Although these approaches provide broad spatial coverage, they often lack real-time, person-centric resolution and raise privacy and computational concerns. By deriving vegetation indices directly from personal-level spectral reflectance, SpectraVita enables real-time, privacy-preserving identification of vegetated, non-vegetated and indoor scenes.

Sensor-derived NDVI emerged as the dominant predictor across classification tasks, with mean absolute β-weight 1.7-fold higher than the second most influential feature. Image-segmentation validation (r = 0.77 between NDVI and vegetation cover) confirms that NDVI reflects actual green space exposure rather than proxy estimates from satellite or location data. This result aligns with established vegetation remote sensing principles and extends them to wearable, individual-level exposure assessment. It also improves interpretability (the decision boundary maps onto an established optical index rather than a black-box embedding) and enhances portability: any wearable with red and near-infrared channels can compute NDVI and implement a comparable, lightweight classifier.

Our multi-stage validation supports translational readiness. Independent scenario-shift testing (100% accuracy) demonstrates generalizability beyond the training distribution. Image-based ground truth validation provides an objective reference standard for green space detection, while mixed-lighting and reflection tests characterize operational limits (notable interference at distances ≤50 cm). Most importantly, large-scale deployments across thousands of participants confirm real-world robustness, with nighttime false natural-light detection ≤1% and sensitivity to seasonal variation (natural light exposure declining from 67% in summer to 49% in winter, and green space exposure from 8% to 4%).

Several limitations must be acknowledged. First, spectral characterization is constrained by the sensor’s dynamic range and field of view. Light at mesopic and scotopic levels—particularly nocturnal light exposure^54^—fell below our detection threshold, and small localized sources such as screens are not explicitly captured. Enabling dynamic gain control in the multispectral sensor in auto mode could expand could extend sensitivity into mesopic level. Second, light-scene classification may be affected by glass transmission and atypical spectra, and vegetation detection can vary with solar angle, vegetation type and canopy structure. future iterations could incorporate geometry-aware calibration or adaptive thresholds and additional spectral indices to improve robustness. Third, the 5-minute sampling interval balances battery life and temporal resolution but may miss brief, potentially meaningful exposures.

In summary, these results indicate that a low-cost, lightweight wearable combined with simple classifier can deliver continuous, individualized, privacy-preserving monitoring of both classical photometric quantities (visual and α-opic metrics) and higher-level context (light source and greenness-related environment), supporting large-scale digital health surveillance and clinically deployable, context-aware interventions. The platform’s robustness across diverse conditions, validated through independent testing, image segmentation, and large-scale deployment, establishes its readiness for translational applications in circadian medicine, behavioral interventions, and population health research.

## Methods

### Hardware design

SpectraVita was designed for functioning primarily as a clip-on or pendant but adaptable as a wrist-worn device by replacing the clip with a watch band. The device consists of a silicone cover and a sensor module. The sensor module measures 33 × 33 × 10 mm and weighs 7.9 g, while the silicone cover, sized at 48 × 39 × 24 mm, adds an additional 8.2 g. The sensor module contains a PCB board, various sensors, a battery, and a Bluetooth module, all enclosed within plastic housing. The upper surface of the sensor module is made of transparent acrylic, featuring two windows, each with a diameter of 2.8 mm. The acrylic discs exhibit irradiance transparency exceeding 85% between 390 nm and 1000 nm, remaining above 20% at 320 nm and 1% at 290 nm. Beneath each window lies a diffuser layer, which ensures uniform light distribution, with transparency exceeding 45% between 380 nm and 1000 nm, 20% at 320 nm, and 1% at 290 nm. Under the diffuser layers, one window is equipped with a broadband UV sensor (G365S01M, Hefei Photosensitive Semiconductor Co., Ltd), featuring a peak responsivity at 355 nm and a detection range of 200–370 nm. The other window houses a multi-spectral digital sensor (AS7341, ams-OSRAM AG., Premstaetten, Austria), which includes eight visible light channels with center wavelengths at 415 nm, 445 nm, 480 nm, 515 nm, 555 nm, 590 nm, 630 nm, and 680 nm. The full width at half maximum (FWHM) for these channels ranges from 26 nm to 52 nm. Additionally, the AS7341 sensor includes a near-infrared (NIR) channel centered at 910 nm, a clear channel for unfiltered silicon response, and a flicker detection channel for measuring light flicker frequency. The angular responses of both sensors to incident light were approximately cosine-shaped. The full width at half maximum (FWHM) was about 106° for the multispectral sensor and 113° for the UV sensor (see supplementary methods and figure 1).

Power is supplied by a 210 mAh Lithium coin cell battery (CR2032). Data transmission is enabled by a low-energy Bluetooth module (BK3435, Beken Corporation, Shanghai, China), a highly integrated Bluetooth® Low Energy 5.0 certified device, facilitating efficient and reliable wireless communication. This compact, lightweight, and versatile design ensures the device is suitable for continuous, real-world light exposure monitoring across diverse environments.

### Software and data collection modes

SpectraVita supports two operational modes for data collection: auto mode and manual mode. Auto mode is the default mode. In the auto mode, both sensors collect data continuously at 5-minute intervals, optimizing the balance between sampling frequency and battery life. With this configuration, the device can store up to 6 days of data, while the battery life extends to approximately 30 days. A WeChat Mini Program was developed to facilitate data upload via Bluetooth and enable real-time data visualization. Users are recommended to upload data at least every other day to ensure data integrity. In this mode, the AS7341 sensor operates with an integration time of 180.7 ms and a gain of 4X.

In the manual mode, the integration time of the AS7341 sensor remains fixed at 180.7 ms, but the gain can be adjusted dynamically from 0.5X to 512X based light intensity. Additionally, the flicker detection channel of the AS7341 sensor, which is inactive in auto mode, is activated in manual mode. This mode allows users to perform targeted testing of specific light environments and labelled the testing in the mini program. Upon completion of data collection, key light characteristics, such as illuminance, are displayed in the mini program.

### Calibration

To ensure the accuracy of the multi-spectral sensor (AS7341) within the visible range, a rigorous calibration process was implemented. First, the spectral output of a commercial evaluation kit was aligned with a high-precision spectrometer (HP350, Hangzhou LCE Intelligent Detection Instrument Co.,Ltd, Hangzhou, China) under various lighting conditions using scaling factors. A full-spectrum light source was employed as the reference lighting for calibration. One device was designated as the golden standard, and its outputs under the test lighting were scaled to match the evaluation kit’s output. Subsequently, under identical lighting conditions, the output of each individual device was scaled to match the golden standard, ensuring consistent performance across all devices. This calibration process guarantees the reliability and accuracy of the spectral data collected by the device.

### Validation

We conducted a series of experiments to evaluate the linearity and accuracy of each channel in both the multi-spectral and UV sensors. To assess the linearity and consistency of the multi-spectral sensor, we tested seven devices under reference light intensities ranging from 2 to 10,753 lux. Measurements were performed using a Spectral Flickering Irradiance Meter (SFIM-400, Hangzhou EVERFINE Photo-E-Info Co., Ltd., Zhejiang, China; wavelength range: 380–780 nm), which served as the reference instrument. All experiments were carried out in a black box, with each sensor positioned beneath the light source. The measurements were collected in manual mode. The coefficient of variation (CV) was used to evaluate the consistency between devices for a given channel and illuminance level, calculated as:

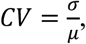

where σ is the standard deviation and μ is the mean of the measurements obtained from the seven devices. A lower CV indicates better inter-device consistency.

To further validate measurement precision for the UV and NIR channels, two light sources were employed: (1) natural sunlight for high-intensity tests, and (2) a UV lamp for low-intensity UV measurements. Sunlight data were collected between 9:00 and 20:00 under clear-sky conditions. During these tests, our device and optical power meters—a UV power meter (HPCS-330UV, Hangzhou HOPOO Optical Color Technology Co., Ltd., Hangzhou, China) and an IR power meter (HPL-220IR2, Hangzhou HOPOO Optical Color Technology Co., Ltd., Hangzhou, China)—were placed side by side on a horizontal, unobstructed surface, facing upward. To ensure comparability, the time offset between measurements from each device was maintained within 3 s (see supplementary results and figure 2).

Finally, to assess the accuracy of spectral features derived, such as illuminance, we reconstructed spectral data from our devices under both natural light and 13 common commercial lamps (including incandescent, fluorescent, and LED). The reconstructed spectral features were then compared with corresponding measurements obtained using a Spectral Irradiance Colorimeter (SPIC-300, Hangzhou EVERFINE Photo-E-Info Co., Ltd., Zhejiang, China; wavelength range: 380–780 nm) under identical conditions. The mean absolute error (MAE) was calculated as the deviation between our device measurements and those from the reference spectrometer:

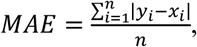

where *y*_*i*_ represents our device’s measurement under condition i, *x*_*i*_ denotes the corresponding spectrometer measurement, and n is the number of test conditions.

### Machine learning for lighting environments classification

We collected SpectraVita data from six primary lighting environments to develop and evaluate our lighting source classification model. These included four indoor settings—LED, fluorescent light (abbreviated as *FL*), incandescent light (*INC*), and indoor sunlight (*SunIn*)—and two outdoor settings—green space (*GS*) and non-green space (*NGS*). To better simulate real-world scenarios, the wearable multispectral sensor was either worn as a pendant and moved naturally within the environment, or fixed at chest level facing forward. For outdoor non-green space environments, long-term continuous monitoring was conducted to capture lighting variations throughout days. Main location includes the rooftop of a 22-floor building (height = 99.1 m) at the West Campus of the University of Science and Technology of China (USTC, 117.3°E, 31. 8°N), Hefei, China—the tallest structure within a 500 m radius—ensuring unobstructed natural light measurements.

Two independent datasets were collected (over 36000 samples): a primary dataset for model training and selection, and a separate validation dataset acquired under completely different scenarios to test model generalizability. Because natural sunlight is highly redundant, primary-dataset daylight samples were further stratified by: (1) cloud cover—sunny (≤50%) vs. rainy (>50%; from https://openweathermap.org); (2) time of day—sunrise/sunset (within 1 h of sunrise or sunset) vs. daytime; and (3) collection location. Data were then down-sampled to obtain approximately balanced sample sizes across these factors. Artificial-lighting measurements with repeated recordings were similarly down-sampled by random selection.

A total of 12 input features were constructed from sensor output. Raw data from channels F1–F8 were normalized by their sum to reduce intensity-related variation. The Clear, NIR, and UV channel values were log-transformed after adding 1 to stabilize variance. In addition, the Normalized Difference Vegetation Index (NDVI) was computed as (NIR-F8)/ (NIR+F8), where NIR and F8 refer to the raw outputs of their respective sensor channels.

Both datasets were first pooled for t-SNE visualization and noise removal to ensure consistent data quality but kept strictly separate for all subsequent modeling. The final primary dataset contained 4309 samples, and the independent validation dataset contained 293 samples (see Supplementary Tables 1 and 2 for details).

Except for semantic segmentation part, all modeling and data analysis were run in MATLAB. For model training and selection, we adopted a stratified, grouped 10-fold cross-validation procedure to ensure robust generalization across lighting conditions. For data organization, measurements from artificial light sources of the same brand were grouped into single sub-categories to minimize intra-class variation. To mitigate class imbalance among sub-categories especially for artificial lighting, we employed the Synthetic Minority Over-sampling Technique (SMOTE) to generate additional synthetic samples until each sub-category contained a comparable number of samples. The total number of generated samples was constrained to no more than three times the original sample size to maintain data diversity and prevent overfitting.

A suite of candidate machine learning algorithms from MATLAB Statistics and Machine Learning Toolbox was evaluated. To compensate for residual class-size differences, sample weights were integrated into the loss function during model training. The final model (a linear support vector machine, SVM) was selected based on a comprehensive evaluation of validation and test performance across 51 random-seed data splits. The primary selection criterion was the highest macro-averaged F1 score, with precision, accuracy, and recall used as secondary criteria.

Note that the datasets used for model development (observations with illuminance >5 lux) were predominantly collected in a manual sampling mode with dynamic gain control, whereas long-term field deployment runs in an automatic mode with a fixed, lower gain. The fixed-gain configuration improves power efficiency and measurement stability for continuous monitoring but reduces low-light sensitivity. To avoid unreliable measurements and keep consistency, model inference in the subsequent analysis was therefore restricted to samples with illuminance >10 lux.

### The contribution of each input variable to machine learning model

The contribution of each input variable was assessed from the coefficients of a linear SVM under K-fold cross-validation. For each fold the model was trained on the training partition only, with all predictors standardized using the training-set mean and standard deviation. We recorded the absolute coefficient vector |β| for each fold and aggregated these values (mean across folds) to obtain a robust importance score per variable. To assess NDVI’s relative importance, predictors were ranked within each fold by |β| (rank 1 = largest), and NDVI’s rank distribution across folds was reported.

### Validation of green-space detection using semantic segmentation

Because an operational definition of green space (e.g., trees, shrubs, grass, dried grassland) is inherently ambiguous, we validated the green-space classification of our final model using image-based semantic segmentation to quantify vegetation in the corresponding scenes. First, we used two synchronized acquisition modules to obtain temporally aligned multimodal data: Wearable multispectral sensing – environmental spectral data were recorded by SpectraVita. Street-view imaging – photographs were captured simultaneously using a GoPro HERO13 (GoPro, Inc., US) equipped with the Ultra-Wide Lens Mod (177° ultra-stabilized field of view; effective visual field = 145° × 113°). In total, 320 paired datasets were collected under diverse configurations and at various times of day. From SpectraVita data, NDVI values and scene categories predicted by the linear Support Vector Machine classifier were extracted.

We then adopted a conservative two-stage procedure of image-based semantic segmentation. First, A DeepLabV3+ network^55^ was trained on the CamVid dataset^56^ using Python, where 32 original category labels were consolidated into 11, and the *“Tree”* label in the new set combined the original *“Tree”* and *“Vegetation Misc”* categories. A ResNet-50 backbone was selected, achieving a mean Intersection over Union (mIoU) of 80.5% on the validation set, a mean mIoU of 74.7% on the test set for the “Tree” category, indicating reasonably high segmentation performance. The corresponding street-view images we collected were resized to meet the DeepLabV3+ input resolution requirements and processed to predict Tree label. Second, human annotators manually removed obvious false positives (e.g., built structures, roads, vehicles and water bodies). Annotators followed a written guideline, and were instructed to only correct clear errors rather than re-define borderline cases. To evaluate the reliability of this manual correction procedure, a randomly selected subset comprising 15% of the scenes (48 images) was independently reviewed and annotated by two annotators. Inter-annotator agreement was assessed based on the resulting vegetation proportion, yielding an intraclass correlation coefficient of 0.98 and a mean difference of 3%. Given the excellent agreement between annotators, the remaining images were subsequently reviewed by the primary annotator.

Finally, the vegetation proportion for each scene image was defined as the proportion of pixels classified as “Tree” relative to the total number of image pixels after removing obvious false positives. These vegetation proportions were then binned into intervals to analyze the probability of scenes being classified as *green space* by SpectraVita across increasing vegetation proportions. An S-shaped (sigmoidal) curve fitting was subsequently applied to model this probability distribution and evaluate the fit quality:

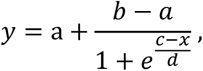

Where a is the y value at the bottom plateau, b is the y value at the top plateau, c is the x value when the response is halfway between bottom and top, 1/d represents the steepness of the curve. The same equation was used in the analysis of Fig 2c-e.

### Validation in a mixed outdoor environment

To examine the diurnal variation in NDVI and the model-derived green-space probability, devices were deployed in one meadow on the west campus of USTC and mounted at chest height, facing green trees located more than 5 m away with 3 approximately orthogonal viewing directions. Data were collected from April to May (6187 samples with illuminance larger than 10 lux in total). Using the geographic coordinates of Hefei and the time of data collection, solar azimuth was calculated according to standard astronomical solar position equations^57^. Device azimuth was measured using a compass.

### Validation in a mixed natural–artificial environment

Measurements were conducted between 7 March and 22 March 2025 in a narrow room on the 13th floor of the USTC building described above, without external artificial light intrusion. The room had a single north-facing window. Four devices were mounted on the long side wall at chest height, facing forward, at distances of 0.5 m, 2.0 m, 3.5 m, and 5.0 m from the window. Two lighting conditions were tested. In the “sunlight only” condition, all artificial light sources were switched off (3778 samples in total). In the “sunlight + LED” condition, LED lights remained on throughout the day (7700 samples).

### Validation in a reflective environment

In a closed room, brightly colored fabric panels (red, green, blue, and white; 65 × 90 cm) were mounted on a wall. Three LED lamps with different correlated color temperatures (4000 K full-spectrum, 5500 K cool, and 3000 K warm), one incandescent lamp, and one fluorescent lamp were each switched on individually from ceiling mounts while all other room lights were off. SpectraVita was mounted at the same height, facing the wall, and positioned seven distances from the wall: 0.25, 0.50, 0.75, 1.00, 1.50, 1.75, and 2.00 m. At each distance under each lighting condition, two replicate measurements were recorded and averaged.

### Validation in a mixed outdoor environment

Four devices were deployed in the center of one meadow on the west campus of USTC and mounted at chest height, facing four orthogonal directions with trees and buildings far away. Data were continuously collected in March 2024 (7 days), May 2025 (7 days) and August 2025 (9 days).

### Human cohort study

To investigate the light that people actually experience in modern daily life, we analyzed data from a cohort study involving school-aged children. The study was conducted in accordance with the ethical guidelines of the First Affiliated Hospital of the University of Science and Technology of China, and written informed consent was obtained from all participants (medical ethics approval number: 2023KY097 and 2024KY157).

Students from eight primary schools of four districts and one county in Hefei, Anhui, China, were invited to participate. Each was instructed to wear SpectraVita in pendant form for at least 14 days per session. The study has been ongoing for multiple years, with students participating in one or two sessions of wearable-sensor data collection. Devices were worn throughout the day, except during sleep and bathing. At night, the device was placed facing upward on the bedside to enable continuous ambient-light measurement. This protocol ensured consistent monitoring of both indoor and outdoor light exposure across daily activities. Each data sample with illuminance larger than 10 lux was classified into environmental scene categories using the trained linear SVM model.

For nighttime analyses (20:00–04:00), we included only participants with more than 4 h of valid predictions. This yielded 218,158 samples from 738 children (8.61 ± 0.35 years old, 368 males) in October–December 2023, 147,260 samples from 570 children (9.16 ± 0.31 years old, 289 males) in May–June 2024, and 199,502 samples from 802 children (8.99 ± 0.94 years old, 409 males) in November 2024–January 2025.

For full time analysis, non-wearing or covered periods were identified when the variance of adjunct sensor signals remained below 10% for more than four consecutive samples. Only participants with ≥ 7 days of valid recordings, and for whom non-wear or covered segments accounted for less than half of the total data, were included in the analysis. Among them, 193 children participated in both sessions of 2024 were included in the full-time seasonal comparison. For each participant, the proportion of predicted labels was calculated for each of the 288 five-minute time bins per day. Daily proportions were then averaged across all subjects to generate the group-level exposure profile.

## Acknowledgements

We thank our youth participants for participating in this study and their parents for supporting. This work would not have been possible without them. Additionally, we are grateful for schools participated in this study, including Hefei Nanmen Primary School, Hefei Nanguohuayuan Primary School, Hefei Normal University Affiliated Xi’an Road School, The Third Primary School Affiliated to Hefei Normal University (Guiyang Road Primary School), Primary School Affiliated to Hefei Normal University and Hefei Tongan Primary School, Hefei 168 Rose Garden School East Campus, and Feidong County Guhe Road Primary School. We acknowledge the support of Hefei Health Commission and Education Commission of Luyang, Binhu, Jingkai and Baohe,distincts. We also thank Zhu yuqiu and the other volunteers for their tremendous efforts in coordination, recruitment, data collection and guidance of study participants. This work was funded by the Science and Technology Innovation 2030 Major Program (2022ZD0210000 to R.L., and 2022ZD0204801 to Y.Z.); the Natural Science Foundation (grant no. 32241010, 32121002, 81925009, and 32430046 to T.X.; 82201230 to R.L.); the CAS Project for Young Scientists in Basic Research (YSBR-013 to T.X., YSBR-041 to R.L.); the Feng Foundation of Biomedical Research to T.X.; and the Human Frontier Science Program (RGY-0090/2014 to T.X.). This work has been supported by the New Cornerstone Science Foundation through the New Cornerstone Investigator Program and the XPLORER PRIZE.

## Author contributions

Conception and design: T.X., and R.L.; Data collection, R.L., Y.H., H.L; Analysis and interpretation, R.L., and Y.H.; writing, R.L., Y.H., and T.X.; supervision, Y.Z and T.X.; funding acquisition, L.R., Y.Z and T.X.

## Competing interests

The authors declare that they have no competing interests.

## Data availability

The datasets generated and/or analyzed during the current study are available from the corresponding author upon reasonable request.

## Code availability

The underlying code for this study is not publicly available but may be made available to qualified researchers on reasonable requests from the corresponding author.

## Supplementary Materials

Supplementary Methods

Supplementary Results

Supplementary Figures

Supplementary Tables

Supplementary References

## Notes

### Competing Interest Statement

The authors have declared no competing interest.

