## Supplementary for "From Spectra to Digital Phenotypes: Wearable Multispectral Sensing for Precision Light and Green Space Exposure"

**This file includes:**

**Supplementary Methods**

**Supplementary Results**

**Supplementary Figures**

**Supplementary Tables**

**Supplementary References**

**Supplementary Methods**

**Measurement of Angular Response for Wearable Light Sensors**

The angular (directional) response of the wearable light sensor was characterized using point sources. The normalized illuminance was measured as a function of the angle of incidence in two orthogonal planes: the sagittal plane and the coronal plane. The measurement was performed inside a darkroom to eliminate ambient light interference. For the multispectral sensor, a single unit was mounted on a flat plate marked with angular scales, and a movable white LED point light source was positioned in front of the device. For each plane (sagittal and coronal), the source was moved along an arc centered on the sensor, maintaining a fixed distance. The angle of incidence varied from -90° (toward the device edge farthest from the sensor) to 90° (toward the device edge closest to the sensor) in 10° increments. For the ultraviolet (UV) sensor, a fixed UV point light source (Thorlabs M365LP1, Thorlabs, Inc., United States) was used. In this configuration, the device was mounted on a rotatable, scale-marked flat plate, with a fixed distance from the source, and the incidence angle was adjusted by rotating the device relative to the light source.

**Extracting common light metrics**

In this study, we extracted several light metrics that are biologically relevant to human visual and non-visual responses. Among them, the α‑opic Equivalent Daylight Illuminance (EDI) serves as a physiologically meaningful metric to quantify the potential of a light source to stimulate the five classes of human photoreceptors in the retina. Unlike conventional illuminance in lux—which reflects only the photopic response of long- and medium-wavelength–sensitive cones (L and M cones)—the α‑opic EDI provides a comprehensive assessment of light’s influence on both visual perception and non-visual processes such as circadian rhythms^1-5^. The prefix “α” corresponds to the five photopigment classes in the human eye: Melanopic (intrinsically photosensitive retinal ganglion cells, ipRGCs), the primary contributor to non-visual effects such as circadian entrainment; Cyanopic (short-wavelength–sensitive cones, S‑cones); Chloropic (medium-wavelength–sensitive cones, M‑cones); Erythropic (long-wavelength–sensitive cones, L‑cones); and Rhodopic (rods, responsible for scotopic or low‑light vision). We computed the α‑opic EDI for each of these photoreceptor types based on the CIE Standard S 026:2018^4^, using th**e** Silent Substitution Toolbox^6^ in Matlab. Briefly, the absolute spectral power distribution of each light condition was weighted by one of the five α‑opic spectral sensitivity functions. The CIE 1964 standard 10‑degree observer was used in all cases, as our focus was on ambient light exposure rather than directed viewing. The age of observers was adjusted to the mean age of our participants using Silent Substitution toolbox. The α‑opic irradiance thus obtained was converted into an illuminance value equivalent to that produced by the CIE Standard Daylight Illuminant D65, representing the same level of photoreceptor activation.

In addition to the α‑opic metrics, we computed conventional photometric and colorimetric measures. We calculated color‑opponent components to represent relative photoreceptor activations. These were defined as the ratios of individual photoreceptor irradiances—L, M, S cones and ipRGC (melanopic)—to the sum of L and M irradiances, denoted as *l*, *m*, *s*, and *i*, respectively. This transformation normalizes cone responses to isolate chromatic mechanisms while discounting luminance variations. We further extracted Correlated Color Temperature (CCT) and tristimulus values (x, y, z) following CIE 1976 Uniform Color Space.

**Supplementary Results**

**Quantification accuracy of SpectraVita**

As illustrated in Supplementary Fig. 2a, sensor outputs from randomly selected seven devices exhibited strong linearity with illuminance across more than four orders of magnitude (2–107553 lux, *n* = 7) under a full-spectrum LED light source. Linear regression analysis yielded coefficients of determination (R²) exceeding 0.99 (mean ± SD: 0.999 ± 0.001), with regression slopes approaching unity (0.99 ± 0.01). The inter-device consistency, evaluated using the coefficient of variation (CV), demonstrated robust reproducibility, with a median CV across all channels (F1–F8, Clear, and NIR) and illuminance levels of 3% ± 5%. These results are consistent with a previous study chosen the same AS7341 sensor^8^.

UV channel performance was further examined under low-intensity irradiation (0.03–1.38 mW/cm²) using a UV lamp. The UV output displayed excellent linearity (R² = 1.00) and inter-device consistency (median CV = 4% ± 5%; *n* = 3). To evaluate performance at higher intensities, the NIR and UV channels were additionally tested under natural sunlight. Both channels maintained strong linear relationships with the spectrometer-derived irradiance values (R² > 0.97), confirming stable operational reliability across dynamic lighting conditions.

Spectrally derived light indices reconstructed from the sensor data also demonstrated close agreement with commercial spectrometer readings. Across diverse artificial and natural lighting conditions (2–43,711 lux; *n* = 16), the reconstructed α‑opic EDI values showed an excellent correlation with reference measurements (R² > 0.99, mean absolute error (MAE) = 0.09 log units, mean slope = 0.97). Except for incandescent illumination, the reconstructed colorimetric indices (e.g., *l*, *s*, *x*, *y*, and *CCT*) also showed high fidelity to spectrometer data (the MAE were 0.0012, 0.0365, 0.0037, 0.0049, 171.78 respectively), demonstrating the system’s strong overall capacity for accurate light environment quantification.

**Supplementary Figures**

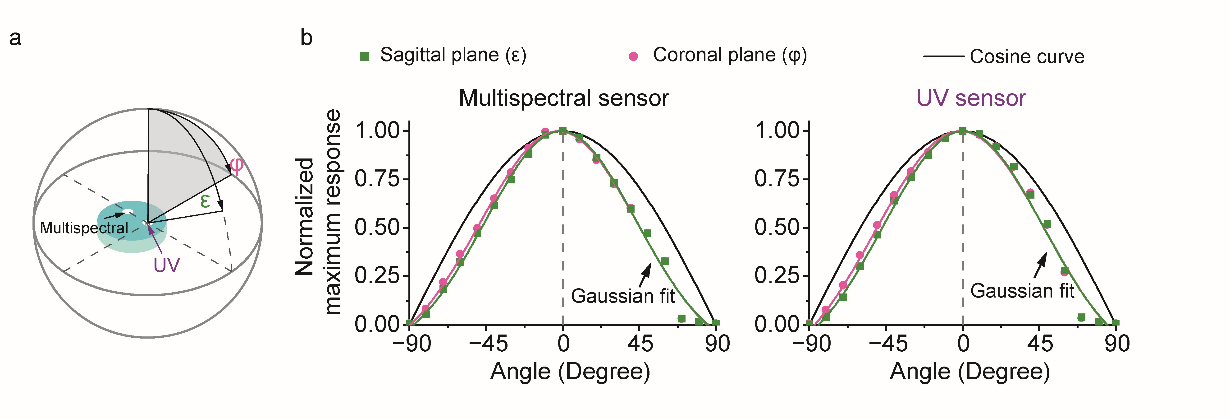

**Supplementary Figure 1.** The directional response of the device. a) Schematic of the SpectraVita sensor module. The two transparent sensor windows (multispectral sensor and the ultraviolet (UV) sensor) are positioned symmetrically along the vertical centerline of the device. Directional response was measured with each sensor centered on the rotation axis. b) Normalized sagittal‑plane (ε, green squares) and coronal‑plane (φ, magenta squares) responses of a single device for the multispectral and UV sensors, measured over incidence angles from −90° (toward the device edge farthest from the sensor) to 90° (toward the device edge closest to the sensor). Solid magenta and green curves show Gaussian fits to the sagittal‑ and coronal‑plane responses, respectively (R² > 0.98). The mean full width at half maximum (FWHM) is approximately 106° for the multispectral sensor and 113° for the UV sensor across both planes. The black curve indicates the ideal cosine response.

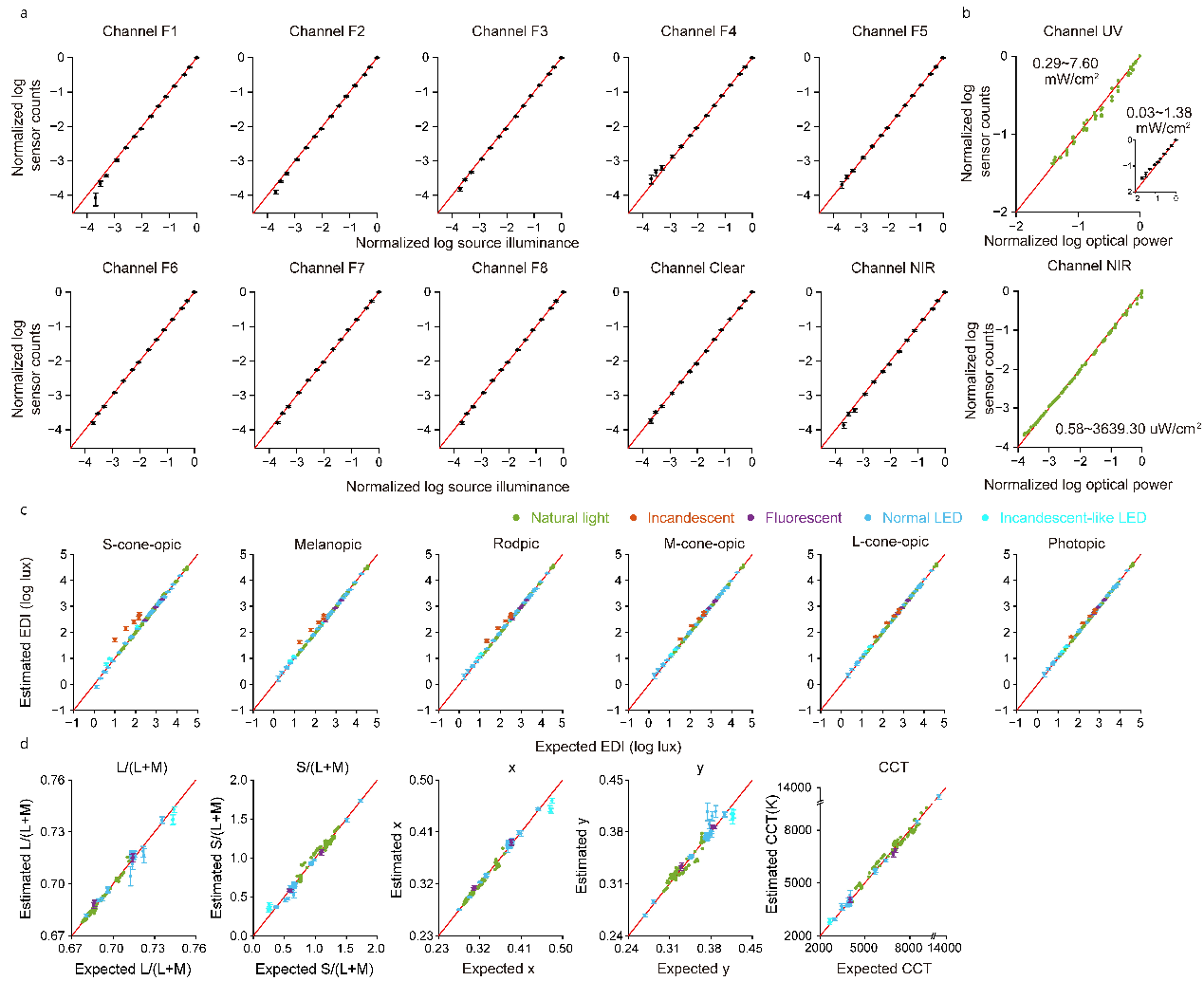
 **Supplementary Figure 2**. a) The linearity and consistency of each visible and NIR channel under a reference LED. The red line represents x = y. b) The linearity of UV and NIR channel under natural light and UV lamp. c) The accuracy of α‑opic EDI compared to a reference spectrometer. d) The accuracy of colorimetric indices compared to a reference spectrometer. Error bars indicate the ±1 SD.

**Supplementary Tables**

**Supplementary Table 1**. The information of the primary dataset

| Light source  (No. of source types) | Location  (No. of location categories) | Sampling period | Data counts (sunny: rainy) |
| --- | --- | --- | --- |
| Sunlight, NGS | Rooftop (2) | 2024.4-2025.1 | 280 (140:140) |
| Sunlight, NGS | Square (8) | 2024.3-2025.6 | 368 (125:243) |
| Sunlight, NGS | Street (4) | 2025.1-2025.3 | 99 (78:21) |
| Sunlight, GS | Tree canopy (17) | 2024.5-2025.8 | 1353 (768:562) |
| Sunlight, SunIn | Room with window (20) | 2024.3-2025.2 | 1313 (842:471) |
| INC (10) | Room without sunlight (7) | - | 250 |
| LED (22) | Room without sunlight (13) | - | 426 |
| FL (10) | Room without sunlight (8) | - | 220 |

**Supplementary Table 2**. The information of the independent scenario-shift validation dataset

| Light source  (No. of source types) | Location  (No. of location categories) | Sampling period | Data counts (sunny: rainy) |
| --- | --- | --- | --- |
| Sunlight, NGS | Square (5) | 2024.3-2025.6 | 42 (15:27) |
| Sunlight, NGS | Street (2) | 2025.1-2025.3 | 14 (6:8) |
| Sunlight, GS | Tree canopy (7) | 2024.3-2025.2 | 48 (32:16) |
| Sunlight, SunIn | Room with window (9) | 2024.5-2025.2 | 47 (33:14) |
| INC (3) | Room without sunlight (2) | - | 33 |
| LED (10) | Room without sunlight (8) | - | 73 |
| FL (5) | Room without sunlight (3) | - | 36 |
